# Sequencing and *de novo* assembly reveals genomic variations associated with differential responses of *Candida albicans* ATCC 10231 towards fluconazole, pH and non-invasive growth

**DOI:** 10.1101/302943

**Authors:** Gajanan Zore, Archana Thakre, Rajendra Patil, Chaithra Pradeep, Bipin Balan

## Abstract

The whole genome sequencing generated 8.09 million paired-end reads, which were assembled into 2,262 scaffolds totaling 17,113,050 bp in length. We predict 7,654 coding regions and 7,647 genes, and annotated the coding regions with gene ontology terms based on similarity to other annotated genomes. Genome comparisons revealed variations (including SNP and indels in genes involved in pH response, fluconazole resistance and invasive growth) compared to *C. albicans* SC5314 and WO1.

---

*Candida* species are important opportunistic pathogens, especially of immunocompromised individuals, where they exhibit high mortality rates (40-75%)^1–6^. The ability of these species to change morphophysiology according to microenvironment contributes to their success as opportunistic pathogens^1–4^. Considering its clinical and economic impact, the Centers for Disease Control and Prevention (CDC), in consultation with experts from NIH and the US-FDA, put *Candida* in the list of organisms that pose a serious antibiotic resistance threat^5^.

*C. albicans* ATCC 10231(Robin) Berkhout was originally isolated from a man with bronchomycosis is included in the *Candida albicans* Drug Resistance Panel (ATCC®MP8™) recommended by the ATCC for use as a control in ASTM Standard Test Method E9799 (ATCC®MP8™)^1–7^. In addition to FLC resistance, it also exhibits traits like pH non-responsiveness and non-invasive growth^8–10^. Before sequencing, differential responses viz. fluconazole resistance, pH non-response and non-invasive growth was confirmed in our laboratory^7–10^. To understand the molecular basis of these traits, we have sequenced the whole genome, developed a *de novo* assembly and compared it with the reference genomes of *C. albicans* SC5314 and WO1. *Candida albicans* (MTCC227), a type strain of *C. albicans* ATCC10231, was collected from Microbial Type Culture Collection (MTCC), Institute of Microbial Technology, Chandigarh (India)^1^. High-quality genomic DNA was extracted by lysing *C. albicans* cells grown in yeast extract peptone dextrose broth for overnight at 28°C with lyticase and then alkaline lysis method^11^. An intact band on the agarose gel and 1.82 ratio of OD at 260/280 confirmed the quality of genomic DNA preparation. Concentration assessed using nano drop was found to be 141.9 ng/micro L. A library of 8.09 million short insert paired end (2×100) reads of 1.6 Gb total size, with an average insert size of 320 bp was generated using Illumina HiSeq2500 platform by following a standard Illumina protocol (Illumina Inc., Cat. # PE-930-1001) as per manufacturer’s instructions^12–15^ (Table 1, Data Citation 1, 2). Read quality was assessed using base quality score distribution, sequence quality score distribution, average base content per read, GC distribution in the reads, PCR amplification issue, check for over-represented sequences and adapter trimming to exclude low-quality sequence reads (the FastQC version v0.10.1)^12–15^. The numbers of bases trimmed from the 5’ and 3’ end of Read1(R1) were 9 and 5 respectively; while that of Read2 (R2) at 5’ and 3’ end were 10 and 4 respectively^12–15^. The adapter sequences were removed using cutadapt (Version-1.8.1)^12^ *De novo* assembly of raw reads was performed using MaSuRCA (Version-3.1.3) as per Zimin et al. (2013); while processed reads were assembled using SOAPdenovo2 and SPAdes as mentioned in Luo, et al. (2012) and Bankevich et al. (2012) respectively^18–20^. Paired-end reads were assembled into 2,262 scaffolds of an average length of 7,565.45 bp and a total length of 17,113,050 bp using MaSuRCA (Table 1, Data Citation 1, 2; Table 2). While using SPAdes (version 3.6.1), we could assemble paired end reads in to 6339 contigs with average length of 2421.19 bp and total length of 15347969 bp and that of using SOAPdenovo263mer were 28492 contigs with average length of 667.44 bp and total length of 19016774 bp (Table 2). We observed that MaSuRCA performed better than other assemblers for the sample sequenced in this study and it could be due to the intrinsic characteristic of two algorithm (DBG and OLC) merged together^16^. MaSuRCA computes an optimal k-mer size of 71 based on the read data and GC content^16^. To validate the assembly further, high-quality reads were aligned to MaSuRCA assembled genome using BWA, which resulted in 83.57% alignment^18–20^.

**Table 1.**
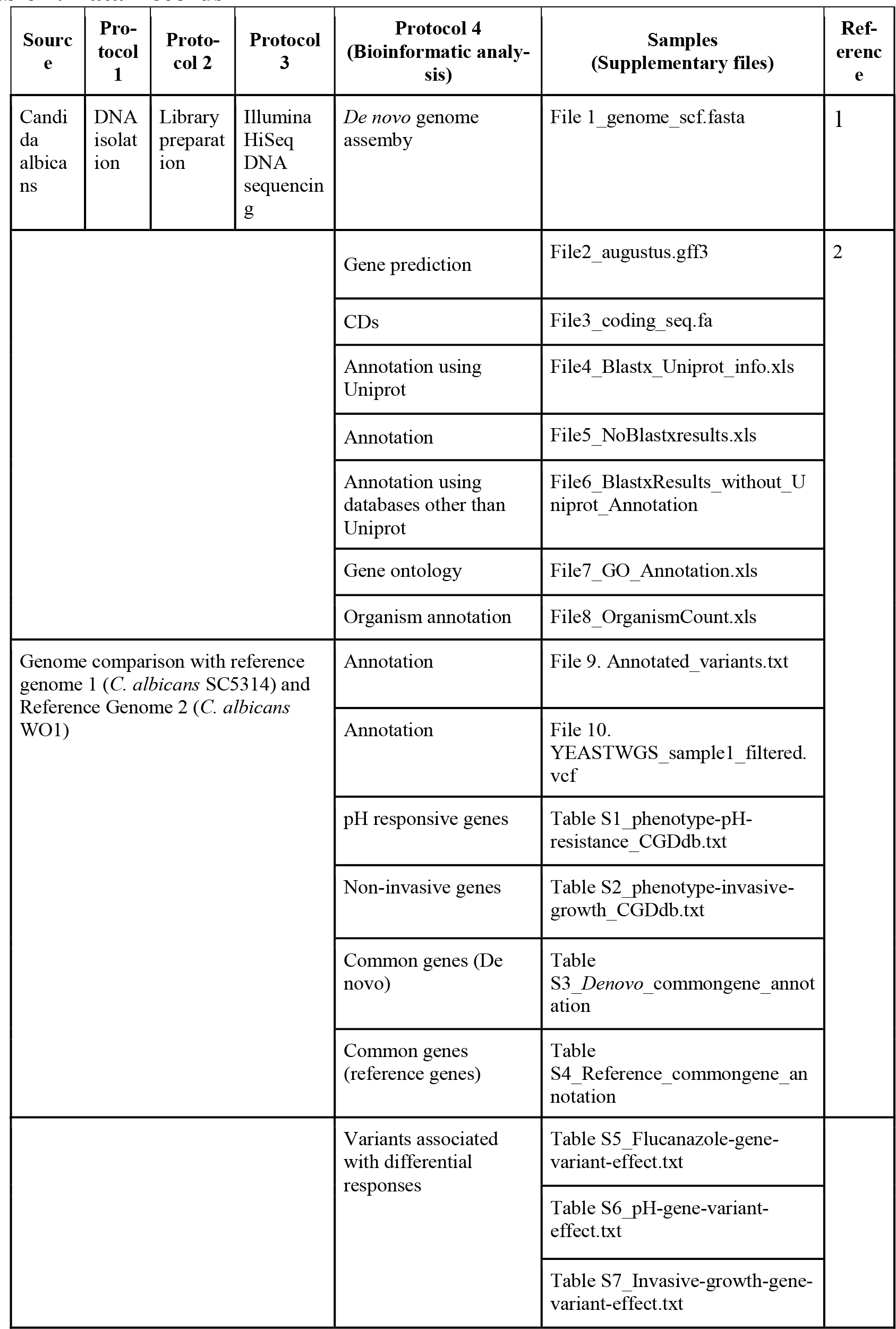
Data Records.

**Table 2.** Genome assembly using different tools.

| Assembler | Contig length (bp) |  |  | N50 | Contigs |  |
| --- | --- | --- | --- | --- | --- | --- |
|  | Minimum | Maximum | Mean |  | Number | Total length (bp) |
| **Spades** | 80 | 110945 | 2421.19 | 8971 | 6339 | 15347969 |
| **Masurca** | 300 | 231110 | 7565.45 | 28273 | 2262 | 17113050 |
| **SOAP** | 100 | 187314 | 667.44 | 4567 | 28492 | 19016774 |

We predict a total of 7,654 coding regions (CDSs) and 7,647 genes from the scaffolds using AUGUSTUS (version 2.5.5) program as per Stanke et al. (2004)^21^.(Table 1, File 2-3, Data Citation 1). Predicted genes were annotated using Perl scripts program using NCBI database and BLASTX program and annotated according to organisms, genes and proteins to the matched genes, gene ontology, and pathways^22–24^.BLASTX hits from the CDS identified 32 organisms (Fig. 1; Table 1, Data Citation 1). The top hit was found to be *C. albicans*, and 99 percent of the genes identified were homologous to *C. albicans* (Fig. 2; File 8, Data Citation 1). Functional annotation terms were predicted for the CDSs from the biological process, molecular function and cellular component branches of the Gene Ontology (GO) (Fig. 3; File 7, Data Citation 1).

**Figure 1:**
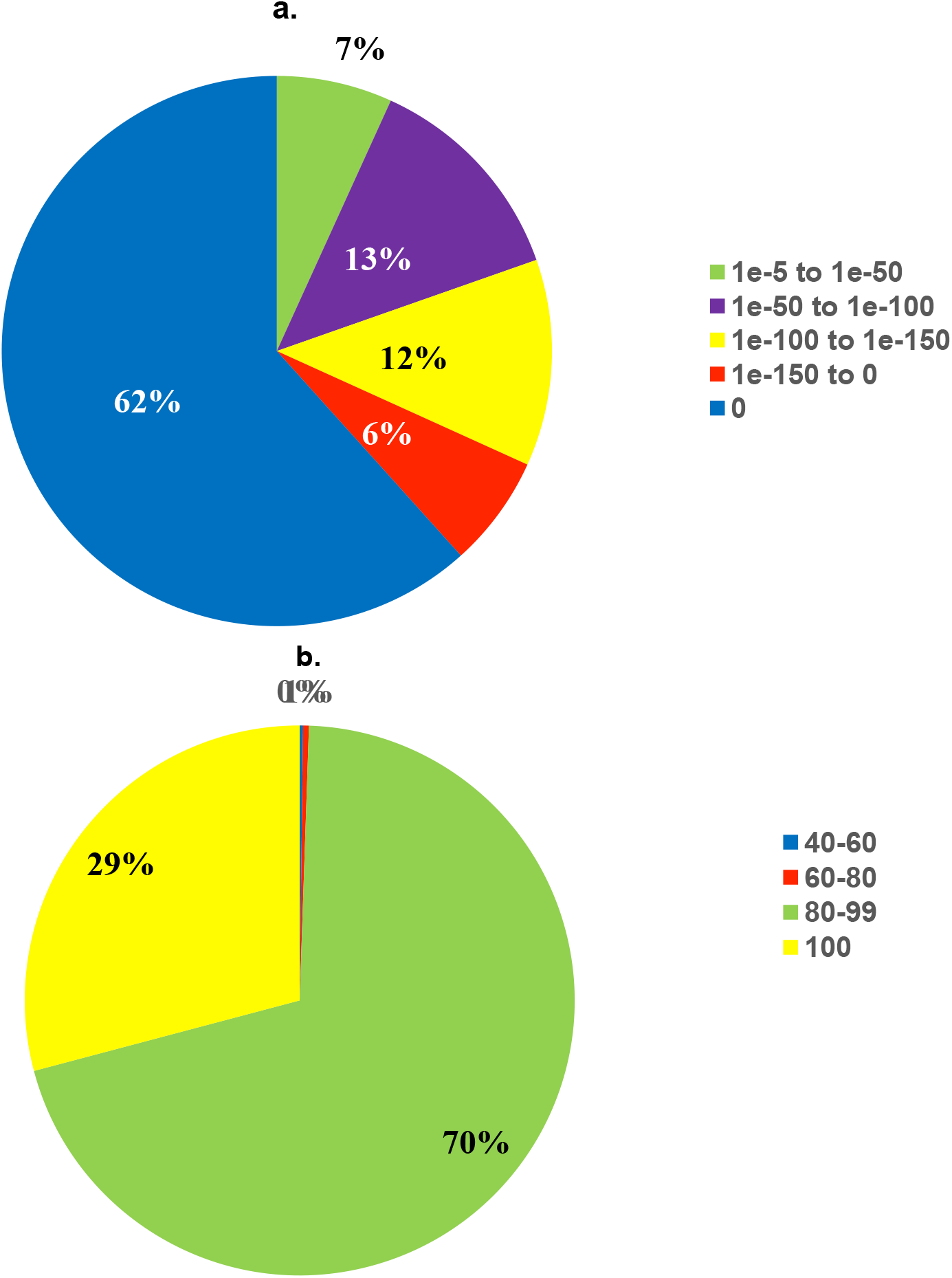
BLASTX E-value distribution and BLASTX similarity score distribution (a) Around 93% of the CDS found using BLASTX have a confidence level of at least 1e-50, which indicates high protein conservation. (b) 99.80% of the predicted CDS found using BLASTX have a similarity of more than 60% at the protein level with the existing proteins at NCBI database.

**Figure 2.**
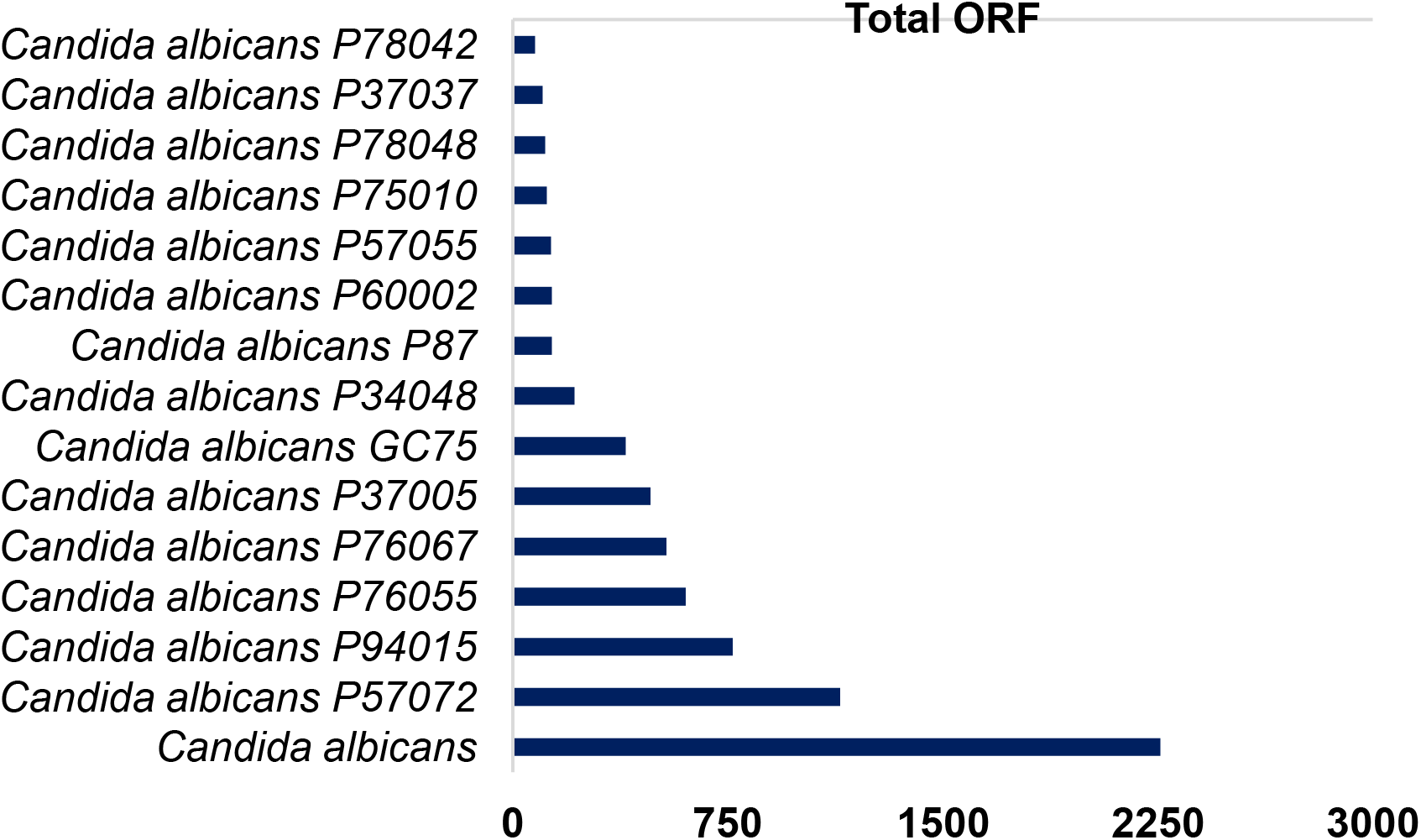
Organism annotation. Top 15 BLASTX organism hits with corresponding ORF count. Among the total 32 organisms found in annotation, the top hit was found to be *C. albicans*.

**Figure 3:**
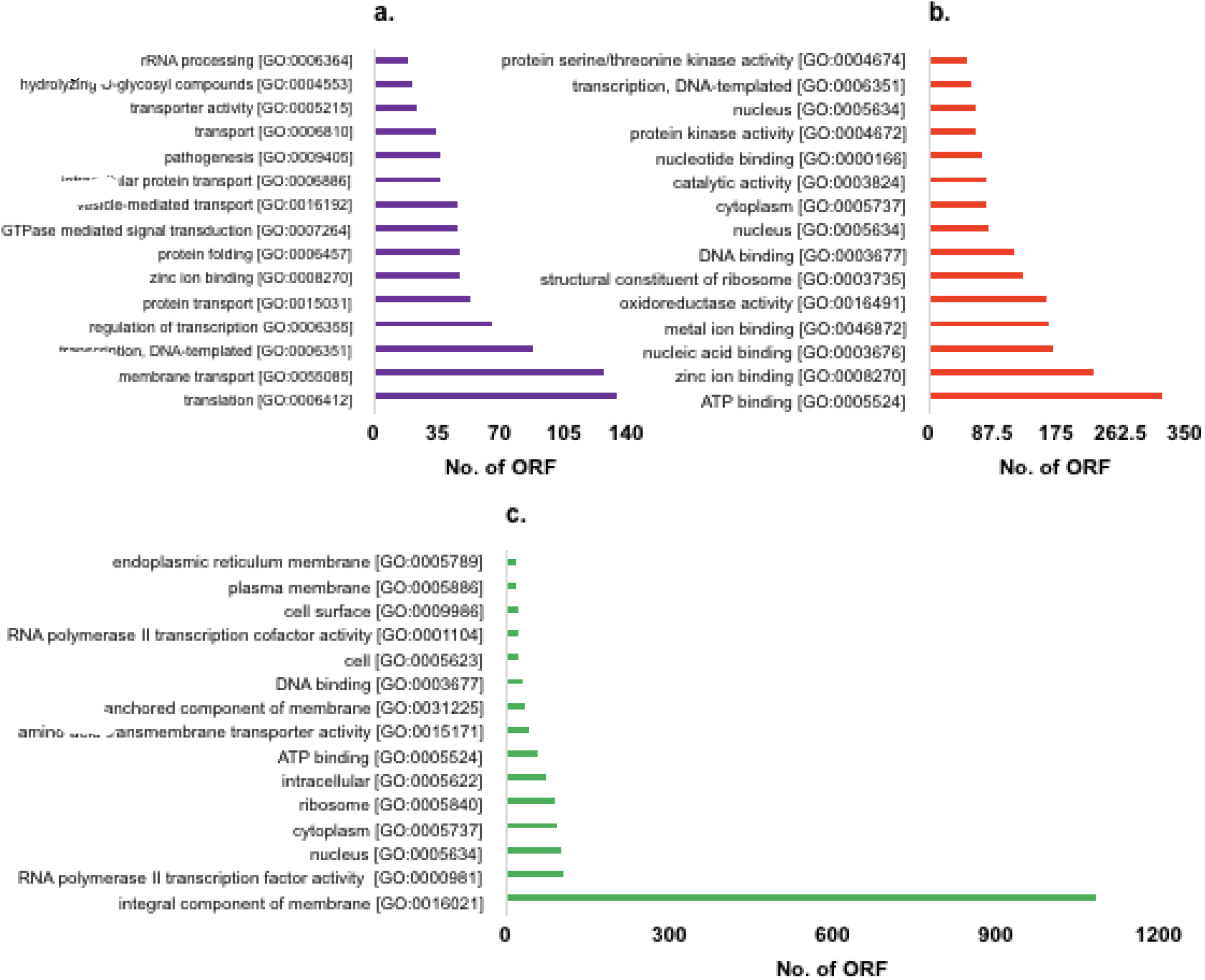
GO annotation according to biological processes, molecular function and cellular component. The top 15 gene ontology (GO) terms for predicted CDS identified in ***(a)*** biological process, ***(b)*** molecular function and ***(c)*** cellular component category.

The variant calling performed for the test genome against the reference genomes of *C. albicans* as well as our *de novo* assembled genome using SAMtools mpileup program revealed global variation, i. e. 35,709 and 94,627 variants compared to *C. albicans* SC5314 and WO1 respectively (Table 3–7)^22–25^. The identified variants were filtered using the cutoffs (read depth ≥= 10 and quality score ≥= 50) ^25^. The details of variants are provided in the table 6. The effect of the identified variants was predicted using SnpEff tool (version 4.3i) against the database created using the assembled genome and its annotation file^25^ The variants identified overall suggest that *C. albicans* ATCC10231 is closer to SC5314 than WO1 (Table 3–7, Fig. 4).

**Figure 4.**
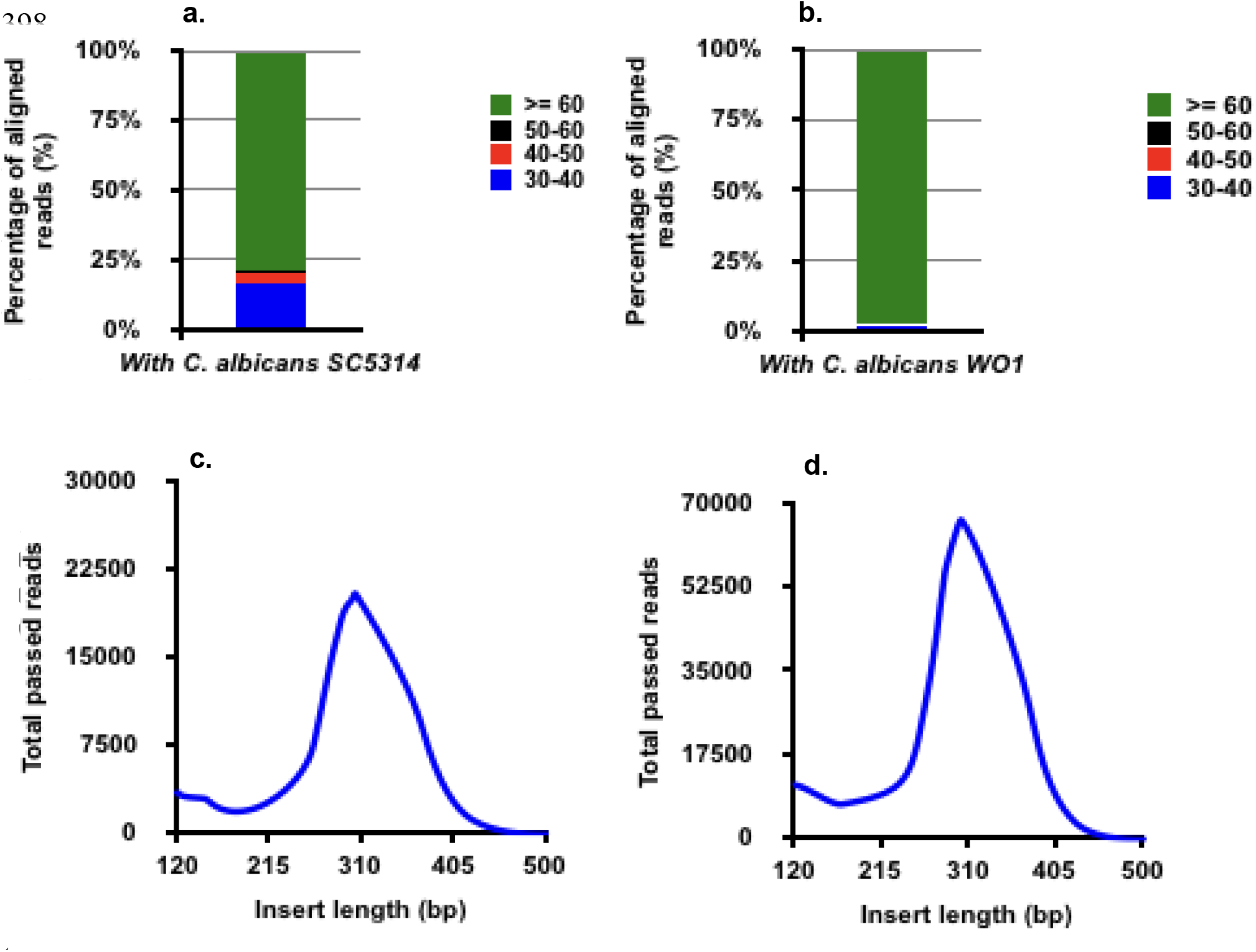
Mapping quality (MAPQ) and Insert size distribution of the aligned reads with *C. albicans* SC531 and *C. albicans* WO1) ***(4a)*** 14. 2 million (~88%) of the total pre-processed reads were mapped to the *C. albicans* strain SC5314 (RG 1) with an average mapping quality of 45.24 and ***(4b)*** 11. 7 million (~72%) of the total pre-processed reads were mapped to the *C. albicans* strain WO1 (RG 2) with an average mapping quality of 54.31. The insert size distribution of the reads with respect to ***(4c)** C. albicans* strain SC5314 (RG 1) and ***(4d)** C. albicans* strain WO1 (RG 2). The X-axis (of c,d), denotes the insert size (in bp) and Y-axis denotes the number of passed reads. The insert size distribution shows a peak at 300bp.

**Table 3.** Quality score distribution.

| Score | SNP | Indel | SNP | Indel |
| --- | --- | --- | --- | --- |
| 50-60 | 2024 | 198 | 2297 | 528 |
| 60-70 | 1926 | 181 | 2762 | 432 |
| 70-80 | 1960 | 158 | 3105 | 392 |
| 80-90 | 1945 | 128 | 3340 | 339 |
| 90-100 | 1984 | 139 | 3645 | 358 |
| >=100 | 22964 | 2102 | 69941 | 7488 |

**Table 4.** Annotation of variants.

| REGION | SNP | INDEL | SNP | INDEL |
| --- | --- | --- | --- | --- |
| Intergenic | 17,052 | 2,108 | 43,979 | 7,166 |
| InsideGene | 15,751 | 798 | 41,111 | 2,371 |
| Exonic | 15,751 | 798 | 40,841 | 2,343 |
| Coding Region | 15,505 | 789 | 40,805 | 2,336 |
| NonCoding Gene | 47 | 3 | 30 | 6 |
| PtCodingGene ncRNA | 199 | 6 | 6 | 1 |
| Intronic | NA | NA | 270 | 28 |
| 5' splice site | NA | NA | 3 | 0 |
| Other | NA | NA | 267 | 28 |

**Table 5.**
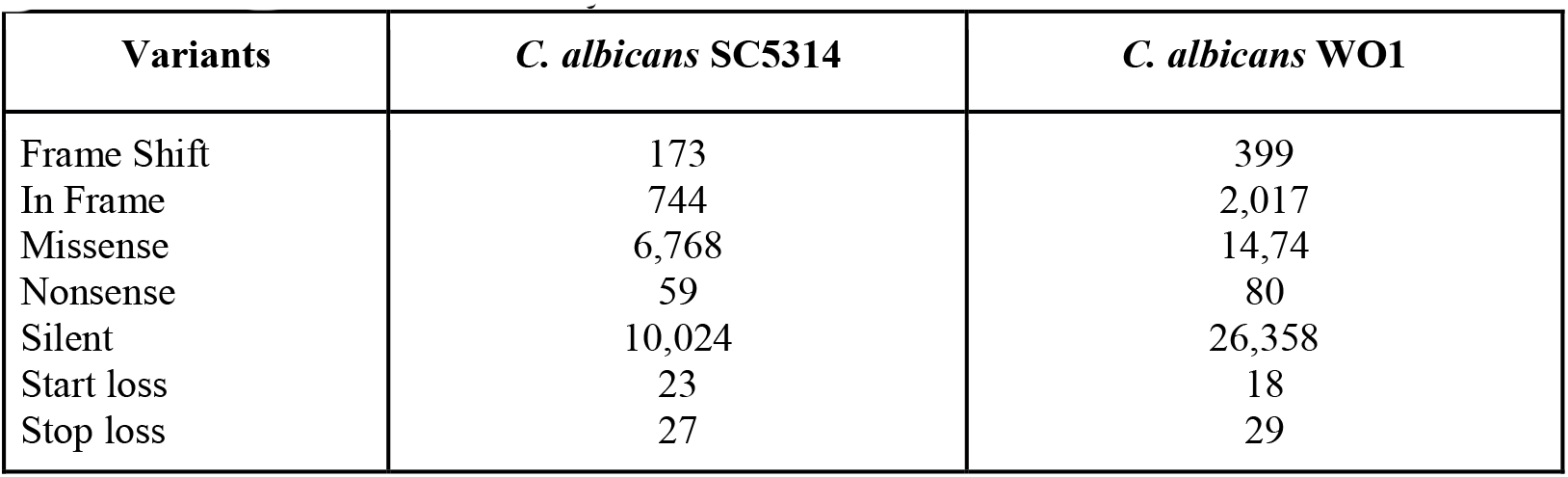
Variant class summary.

| Variants | *C. albicans* SC5314 | *C. albicans* WO1 |
| --- | --- | --- |
| Frame Shift | 173 | 399 |
| In Frame | 744 | 2,017 |
| Missense | 6,768 | 14,74 |
| Nonsense | 59 | 80 |
| Silent | 10,024 | 26,358 |
| Start loss | 23 | 18 |
| Stop loss | 27 | 29 |

**Table 6.** Identified variants.

| Variants | *C. albicans* SC5314 | *C. albicans* WO1 |
| --- | --- | --- |
| Total Variants | 72,036 | 1,11,011 |
| Passed Variants | 35,709 | 94,627 |
| Total SNPs | 32803 (91.86%) | 85090 (89.92%) |
| Total inDels | 2906 (8.14%) | 9537 (10.08%) |
| Total Homozygous SNP | 21606 (65.86%) | 41703 (49.01%) |
| Total Heterozygous SNP | 11197 (34.13%) | 43387 (50.99%) |
| Total transition type SNP | 23324 (71.1%) | 62049 (72.92%) |
| Total Transversion type snp | 9479 (28.89%) | 23041 (27.08%) |
| Ts/Tv | 2.46 | 2.69 |

**Table 7.** Read depth distribution of the identified variants with *C. albicans* SC5314 and WO1.

| Read Depth | *C. albicans* SC5314 |  | *C. albicans* WO1 |  |
| --- | --- | --- | --- | --- |
|  | Total SNPs | Total INDELS | Total SNPs | Total INDELS |
| 0-5 | 0 | 0 | 0 | 0 |
| 5-10 | 0 | 0 | 0 | 0 |
| 10-15 | 2148 | 137 | 712 | 162 |
| 15-20 | 2894 | 344 | 1700 | 453 |
| 20-25 | 2937 | 360 | 2680 | 720 |
| 25-30 | 2719 | 339 | 3774 | 824 |
| 30-50 | 9072 | 971 | 23302 | 3547 |
| >=50 | 13033 | 755 | 52922 | 3831 |

Polymorphic sites within the assembled genome were identified by mapping the reads to reference genomes of C. *albicans* SC5314 (Data Citation 1). Nine genes (viz.VPS36, RIM21, RIM9, CSR1, PHR1, RIM13, vps28, SIT4,CCC1/4) involved in pH response, seven (viz. ERG6, ERG11, CDR3, POR1, CDR4, NDT80, HOG1) in fluconazole resistance and forty (including ENO1, PHR1, CDC10, RSP5, SET1, GPR1, SIT4 and ASH1) in invasive growth showed significant variation (Table 8)^4–6,8–10^. These variants could be responsible for pH non-responsiveness, fluconazole resistance and non-invasive growth in *C. albicans* ATCC10231 (Table 7, Supplementary Table S1-S7, Data citation 1)^26–39^.

**Table 8.**
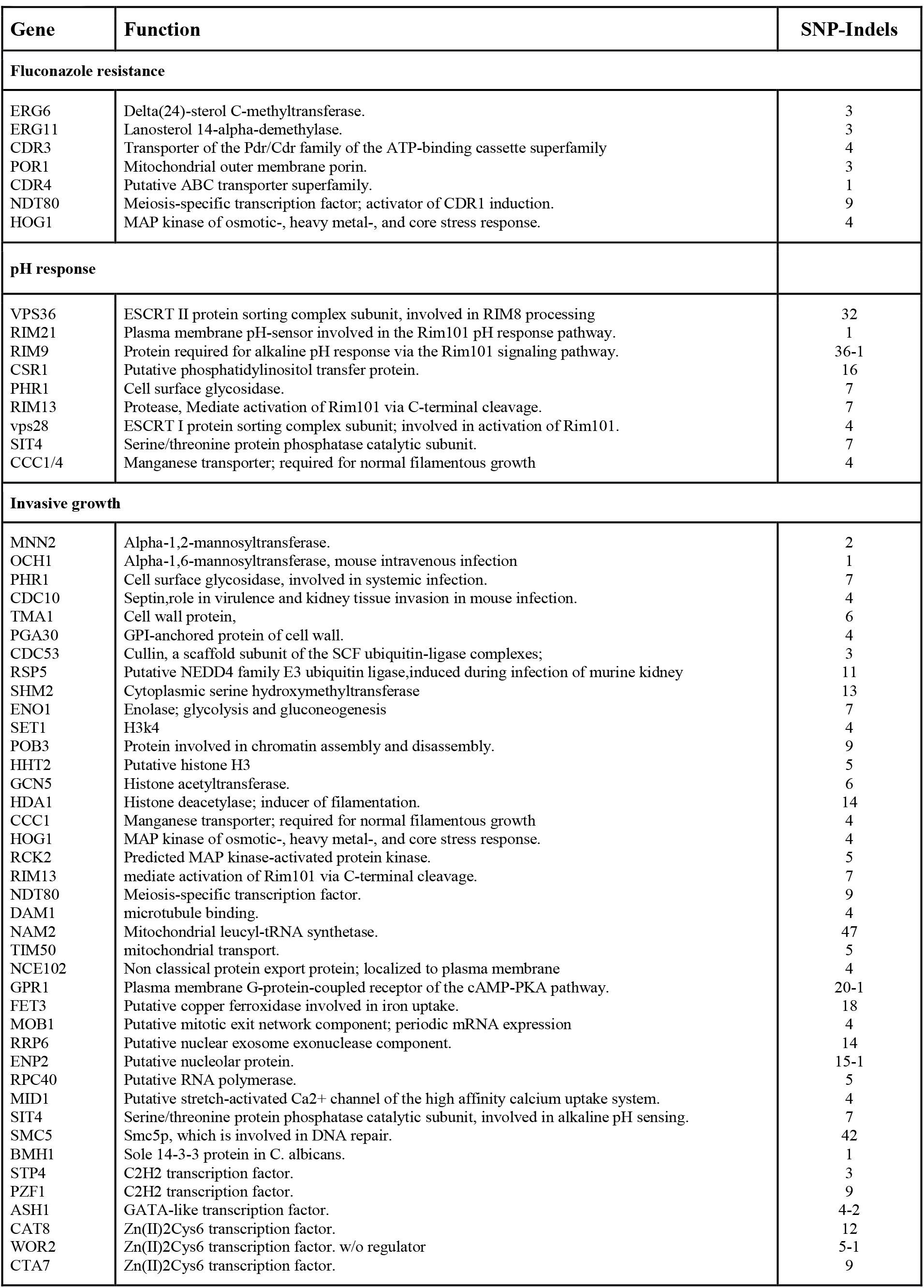
Variants associated with fluconazole resistance, pH response and noninvasive growth in *Candida albicans* ATCC10231.

| Gene | Function | SNP-Indels |
| --- | --- | --- |
| **Fluconazole resistance** |  |  |
| ERG6 | Delta(24)-sterol C-methyltransferase. | 3 |
| ERG11 | Lanosterol 14-alpha-demethylase. | 3 |
| CDR3 | Transporter of the Pdr/Cdr family of the ATP-binding cassette superfamily | 4 |
| POR1 | Mitochondrial outer membrane porin. | 3 |
| CDR4 | Putative ABC transporter superfamily. | 1 |
| NDT80 | Meiosis-specific transcription factor; activator of CDR1 induction. | 9 |
| HOG1 | MAP kinase of osmotic-, heavy metal-, and core stress response. | 4 |
| **pH response** |  |  |
| VPS36 | ESCRT II protein sorting complex subunit, involved in RIM8 processing | 32 |
| RIM21 | Plasma membrane pH-sensor involved in the Rim101 pH response pathway. | 1 |
| RIM9 | Protein required for alkaline pH response via the Rim101 signaling pathway. | 36-1 |
| CSR1 | Putative phosphatidylinositol transfer protein. | 16 |
| PHR1 | Cell surface glycosidase. | 7 |
| RIM13 | Protease, Mediate activation of Rim101 via C-terminal cleavage. | 7 |
| vps28 | ESCRT I protein sorting complex subunit; involved in activation of Rim101. | 4 |
| SIT4 | Serine/threonine protein phosphatase catalytic subunit. | 7 |
| CCC1/4 | Manganese transporter; required for normal filamentous growth | 4 |
| **Invasive growth** |  |  |
| MNN2 | Alpha-1,2-mannosyltransferase. | 2 |
| OCH1 | Alpha-1,6-mannosyltransferase, mouse intravenous infection | 1 |
| PHR1 | Cell surface glycosidase, involved in systemic infection. | 7 |
| CDC10 | Septin, role in virulence and kidney tissue invasion in mouse infection. | 4 |
| TMA1 | Cell wall protein, | 6 |
| PGA30 | GPI-anchored protein of cell wall. | 4 |
| CDC53 | Cullin, a scaffold subunit of the SCF ubiquitin-ligase complexes; | 3 |
| RSP5 | Putative NEDD4 family E3 ubiquitin ligase, induced during infection of murine kidney | 11 |
| SHM2 | Cytoplasmic serine hydroxymethyltransferase | 13 |
| ENO1 | Enolase; glycolysis and gluconeogenesis | 7 |
| SET1 | H3k4 | 4 |
| POB3 | Protein involved in chromatin assembly and disassembly. | 9 |
| HHT2 | Putative histone H3 | 5 |
| GCN5 | Histone acetyltransferase. | 6 |
| HDA1 | Histone deacetylase; inducer of filamentation. | 14 |
| CCC1 | Manganese transporter; required for normal filamentous growth | 4 |
| HOG1 | MAP kinase of osmotic-, heavy metal-, and core stress response. | 4 |
| RCK2 | Predicted MAP kinase-activated protein kinase. | 5 |
| RIM13 | mediate activation of Rim101 via C-terminal cleavage. | 7 |
| NDT80 | Meiosis-specific transcription factor. | 9 |
| DAM1 | microtubule binding. | 4 |
| NAM2 | Mitochondrial leucyl-tRNA synthetase. | 47 |
| TIM50 | mitochondrial transport. | 5 |
| NCE102 | Non classical protein export protein; localized to plasma membrane | 4 |
| GPR1 | Plasma membrane G-protein-coupled receptor of the cAMP-PKA pathway. | 20-1 |
| FET3 | Putative copper ferroxidase involved in iron uptake. | 18 |
| MOB1 | Putative mitotic exit network component; periodic mRNA expression | 4 |
| RRP6 | Putative nuclear exosome exonuclease component. | 14 |
| ENP2 | Putative nucleolar protein. | 15-1 |
| RPC40 | Putative RNA polymerase. | 5 |
| MID1 | Putative stretch-activated Ca<sup>2+</sup> channel of the high affinity calcium uptake system. | 4 |
| SIT4 | Serine/threonine protein phosphatase catalytic subunit, involved in alkaline pH sensing. | 7 |
| SMC5 | Smc5p, which is involved in DNA repair. | 42 |
| BMH1 | Sole 14-3-3 protein in *C. albicans*. | 1 |
| STP4 | C2H2 transcription factor. | 3 |
| PZF1 | C2H2 transcription factor. | 9 |
| ASH1 | GATA-like transcription factor. | 4-2 |
| CAT8 | Zn(II)2Cys6 transcription factor. | 12 |
| WOR2 | Zn(II)2Cys6 transcription factor. w/o regulator | 5-1 |
| CTA7 | Zn(II)2Cys6 transcription factor. | 9 |

Neutral pH response is mediated by a seven domain containing transmembrane receptor, DFG1 that along with other proteins including Vps36 (ESCRT II protein sorting complex subunit), Vps28 (ESCRT I protein sorting complex subunit) undergo endocytosis and associate with Endosomal Sorting Complex Required for Transport (ESCRT)^27–29^. Endocytosed Vps36 and Vps28 activate (phosphorylation and ubiquitination) Rim8 (β-arrestin like protein) that in turn interacts and recruit Rim21 (plasma membrane pH-sensor) and Rim101 (zinc finger transcription factor) at ESCRT complex^27–29^. This interaction brings RIM101 in proximity to RIM13 (protease) that cleaves C-terminal peptide and activate short N-terminal, zinc finger transcription factor, Rim101^27–29^. RIM101 recognizes 5’– NCCAAG-3’ sequence and induce expression of alkaline pH responsive genes like PHR1 (cell surface glycosidase), Rim101 etc., under neutral pH^27–29^. The genes of *C. albicans* ATCC10231 involved in this process was found to exhibit the significant variation (SNP/indel) viz. VPS36 (32), VPS28 (4), RIM9 (36),13 (7) and 21 (1), CSR1 (16), PHR1 (7) and SIT4 (7). These variations could be associated with pH nonresponsiveness of *C. albicans* ATCC10231, as mutants were reported to affect alkaline pH induced hyphae formation and thus virulence^27–29^.

Activation of drug efflux pumps (MDR, CDR), modification and or over expression of drug target are the major mechanisms that confer drug resistance in both pro and eukaryotic organisms/cells^26^. Our data suggest that FLC resistance in *C. albicans* ATCC10231 could be due to either modification in 14-alpha Demethylase (FLC target encoded by ERG11) and or drug efflux pumps^26^. As seven genes involved in ergosterol biosynthesis (ERG11, ERG6), drug efflux and regulator (CDR3, CDR4, NDT80), stress response (HOG1) and a mitochondrial membrane transporter (POR1) exhibit variations (SNP/Indel)^26^. However, it was reported that neither mutations nor over expression of both CDR3 and CDR4 modulate FLC susceptibility, indicating that modified ERG11 could be conferring FLC resistance in *C. albicans* ATCC10231.

In addition to these, forty genes regulating the invasive growth of *C. albicans* ATCC 10231 exhibit significant variations (SNP/Indel) (Table 7, Supplementary Table S5-7, Data citation 1). SNPs were ranged from 1-47 i.e. maximum was observed in Nam2 (47) followed by SMC5 (42) and RIM21 (36) (Table 7, Supplementary Table S5-7, Data citation 1). Genes like ENO1, PHR1, CDC10, RSP5, SET1, GPR1, SIT4 and ASH1 reported to be essential for tissue invasion exhibit 7, 7, 4, 11, 4, 20, 7, 4 SNPs respectively (GPR1 and ASH1 exhibit 1 and 2 indels respectively) (Table 7, Supplementary Table S5-7, Data citation 1)^30–39^. Eno1 is a cell surface, plasminogen binding protein essential for tissue invasion^37^. PHR1 codes for a glucosidase (involved in beta 1,3 glucan processing) is also required for maintaining hyphal morphology, adhesion (by regulating expression of Hwp1 and ECE1) and thus tissue invasion^27^. A septin encoded by CDC10 is essential for invasive growth as mutants though form hyphae failed to penetrate solid surfaces both *in vivo* and *in vitro* and thus affect virulence^30^. NEDD4 family E3 ubiquitin ligase coded by RSP5 is essential in systemic kidney infection in mouse^35^. SET1, an H3K4 methyltransferase is essential for pathogenesis as loss of SET1 affected cell surface chemistry and adherence thus impacting epithelial cell invasion^38^. GPR1 mutants failed to induce invasive growth on solid hypha-inducing media^39^. SIT4 regulates two hyphae specific glucanases involved in cell wall biogenesis as mutants failed to induce invasive growth.

Whole genome sequencing of a strain differentially responsive to pH, invasive growth and anti-fungal agents has provided an insight into the molecular basis of these phenotypes. Thus it may find use as a unique reference database to understand novel mechanisms and survival strategies evolving in eukaryotic organisms in general and *C. albicans* in particular.

## Data Records

The whole genome sequence of *Candida albicans* ATCC10231 is deposited to NCBI Sequence Read Archive (SRA) database as the raw FastQ files, whole genome assembly and annotation under accession number SRP067106 and Figshare. https://doi.org/10.6084/m9.figshare.5349937.v1. Data files available at Figshare are as follows:

## Acknowledgements

Authors are thankful to SERB, Govt. of India, India for financial assistance to GBZ under SERB FTSYS and Savitribai Phule Pune University, Pune (MS) India for financial assistance to RP. Authors are also thankful to Prof. Pandit Vidyasagar, Vice Chancellor, SRTM University, Nanded (MS) India for his encouragement and constant support.

## Author contributions

GBZ conceived, designed the experiment analyzed the data, evaluated the conclusions and wrote the paper. RP performed the experiments. BB and CP performed bioinformatics analysis and evaluated conclusion. The manuscript is read and approved by all the authors before communication.

